# Targeting of immune cells by human adenoviruses: CD46 receptor density influences entry of chimaeric Ad5F35 into natural killer cells

**DOI:** 10.64898/2026.06.08.730800

**Authors:** Tyler Barr, Elif Aktar, Sian L. Drake, Magdalena Karwatka, Erica B. Wilson, Ruth Hughes, G. Eric Blair, Graham P. Cook

**Affiliations:** Leeds Institute of Medical Research, School of Medicine, University of Leeds, UK; School of Molecular and Cellular Biology, Faculty of Biological Sciences, University of Leeds, UK; Bioimaging Facility, Faculty of Biological Sciences, University of Leeds, UK

## Abstract

Most adenovirus (Ad) vectors are based on the genome of human Ad type 5 (Ad5), which targets their entry to cells that express the Coxsackie and Adenovirus Receptor (CAR or CXADR). However, certain human Ads do not use CAR, for example Ad35 interacts with cell-surface CD46 and Ad3 uses desmoglein 2 (DSG2) for cell entry. In this study, a comparison of different Ad receptors using transcriptomic and proteomic databases showed that CD46 is widely expressed across human cells and tissues whereas CAR and DSG2 are more restricted to epithelial cells. We have used a hybrid virus, Ad5F35, that comprises an Ad5 genome in which the Ad5 fibre was replaced with that of Ad35, thus retargeting the virus from CAR- to CD46-expressing cells and enabling transduction of primary human NK and T cells. However, lymphocytes required approximately 10 to 20-fold more Ad5F35 particles per cell (ppc) compared to A549 epithelial cells to achieve a similar level of transduction. Consistent with this, quantitation of the cell-surface density of CD46 molecules revealed approximately 100 CD46 molecules per µm^2^ in primary NK cells compared with approximately 2000 CD46 per µm^2^ in HeLa cells, a 20-fold difference. Cell-surface CD46 density was reduced by approximately 95% by RNA interference in HeLa cells to levels that approximate those found on NK cells. Lower CD46 density reduced transduction by Ad5F35 but this could be compensated for with increased MOI. Our results identify the density of cell surface CD46 as a critical determinant of Ad5F35 transduction and demonstrate that Ad5F35 is an efficient vector for gene delivery in primary human NK cells.

## 1. Introduction

The cell tropism of human adenoviruses (Ad) is primarily determined by interactions between the Ad fibre protein and cell surface attachment molecules [1,2]. Binding of the Ad fibre to a cell surface protein is accompanied by interactions between the Ad penton base and cell-surface integrin molecules, principally of the α_v_ family, such as α_v_β_3_ or α_v_β_5_, with subsequent endocytosis and virus entry [1,2]. Most of the seven human Ad species, A-G, utilise the Coxsackie and Adenovirus receptor (CAR) as a primary entry molecule, this includes the well-studied Ad type 5 (Ad5), the most commonly used Ad in gene delivery and therapy applications [3–6]. The CAR molecule is expressed in a tissue-specific manner and has been studied in greatest detail in epithelial cells, where it plays a role in maintaining tight junctions [6–8]. This restricted expression of CAR may limit the therapeutic applications of Ad5-based vectors [9,10]. In contrast, the species B Ads (for example, Ad11 and Ad35) exploit interactions between their fibre protein and CD46, a widely expressed complement regulatory receptor [11–14]. A third receptor, desmoglein-2 (DSG2), interacts with the fibre protein of certain species B Ads such as Ad3 which, like CAR, has a restricted expression pattern [15].

Targeting of the widely expressed cell-surface molecule CD46 by viruses that contain a species B fibre has the potential to extend the therapeutic use of Ads to many human cell types [16]. This potentially includes the retargeting of cytotoxic T cells and natural killer (NK) cells using chimaeric antigen receptors (CARs) for cancer immunotherapy and optimising the tropism of oncolytic Ads. However, lack of efficient gene delivery to human immune cells remains an obstacle in these approaches. Lentivirus vectors have been used for the *ex vivo* transduction of human lymphocytes, but transduction efficiency is low and extensive *ex vivo* proliferation of transduced cells with cytokines is required to generate sufficient modified cells for therapy [17,18]. Such proliferation is not only time consuming and costly but can also result in functional exhaustion of immune cells, reducing their effectiveness *in vivo* [19]. A higher efficiency transduction system might offer a route towards more efficient gene delivery and therapeutic use of modified immune cells. Furthermore, human lymphocytes (especially NK cells) are refractory to several standard methods of gene manipulation, and this has hampered their study; an efficient, viral based transduction system such as that provided by Ads promises to advance both basic and translational studies of these key immune effector cell types.

In this study, primary human NK cells were transduced with a chimaeric adenovirus vector, Ad5F35, in which the Ad5 fibre was replaced with that of Ad35, targeting the vector to CD46-expressing cells. Quantification of CD46 levels show that lymphocytes have an 10-20-fold lower density of CD46 molecules than epithelial cells, reducing transduction efficiency. We hypothesised that this low CD46 density was compensated for by using a high titre of Ad5F35 for transduction. This is supported by reducing CD46 density on HeLa cells and demonstrating that resultant reduced transduction efficiency can be overcome using a high titre of Ad5F35. Our data further suggests that, while the density of cell-surface CD46 molecules at the cell surface plays an important role in entry of species B Ads, other, as yet unidentified, post-entry mechanisms are also important in Ad transduction and infection of different human cell types.

## 2. Materials and Methods

### Antibodies

The FITC-labelled anti-human CD46 (clone E4.1), APC-labelled anti-human CD56 (clone B159), APC-Cy7 labelled anti-human CD3 (clone SK7) and APC-labelled anti-human CD8 (clone SK1) antibodies were purchased from BD Biosciences (Oxford, UK). PE-Vio770 labelled anti-human CD4 (clone SK3) was from Miltenyi Biotec. Quantum™ Simply Cellular (QSC) beads were from Bangs Laboratories, (Indiana, USA). For blocking experiments, we used rabbit anti-human CD46 polyclonal antibody or rabbit IgG from Santa Cruz.

### Cell Lines

HeLa cells and A549 cells (obtained from the European Collection of Authenticated Cell Cultures (ECACC), Salisbury, UK) were cultured in Dulbecco’s Minimal Eagle Medium (DMEM; Sigma-Aldrich, Dorset, UK) supplemented with 10% Foetal Bovine Serum (FBS; GE Healthcare, Buckinghamshire, UK), 2 mM L-glutamine, 50 units penicillin/mL and 50 µg streptomycin/mL (Sigma-Aldrich). Adherent cells were detached using trypsin/EDTA (Sigma-Aldrich). All cells were cultured at 37°C in a humidified atmosphere of 5% CO_2_.

### Viruses

Ad5-EGFP (Enhanced Green Fluorescent Protein) is a replication-deficient E1- and E3-deleted adenovirus based on Ad5 in which the E1 region was replaced with a cytomegalovirus (CMV) promoter-driven EGFP transgene and Ad5F35-EGFP is identical to Ad5-EGFP virus except that the Ad35 fibre replaced the Ad5 fibre [20]. Viruses were propagated in 911 cells and purified using the freeze-thaw method followed by caesium chloride centrifugation [21]. The number of virus particles (vp) was determined by spectrophotometry, as previously described [22].

### Transduction of A549 and HeLa epithelial cells

Cells (5×10^4^) were plated in six well plates and incubated for 24 h at 37°C in a humidified atmosphere containing 5% CO_2_. The cells were washed with PBS followed by addition of Ad5-EGFP or Ad5F35-EGFP in serum-free DMEM. The cells were incubated for one hour at 37 °C in a humidified atmosphere with 5% CO2, complete DMEM was added and incubated for a further 24h in a humidified atmosphere with 5% CO_2_. The cells were detached by addition of trypsin-EDTA followed by neutralization with “complete” DMEM medium, collected by centrifugation, washed by addition of PBS with centrifugation and resuspension in PBS. Transduced cells were identified by EGFP expression using flow cytometry. For blocking experiments, A549 cells were treated with rabbit anti-human CD46 polyclonal antibody (or rabbit IgG) at a final concentration of 4µg/ml prior to addition of Ad5F35.

### Isolation of Peripheral Blood Mononuclear Cells (PBMC) and purification of NK cells

PBMC were obtained from healthy donor leukocyte apheresis cones supplied by the National Health Service Blood and Transplant unit (NHSBT), as described [23]. Cells were eluted from cones and PBMC isolated by density gradient centrifugation using Lymphoprep^TM^ (STEMCELL Technologies, UK) and cultured in RPMI supplemented with 10% FBS. For NK cell isolation, PBMC were fractionated using indirect immunomagnetic selection (Miltenyi Biotec). The purity of the isolated population was assessed by staining with anti-CD3 and anti-CD56 antibodies and NK cells identified as the CD56+CD3^neg^ population; this method routinely generates populations of NK cells with purity exceeding 95%. For blocking experiments, purified NK cells were treated with rabbit anti-human CD46 polyclonal antibody (or rabbit IgG) at a final concentration of 4µg/ml prior to addition of Ad5F35.

### Transduction of PBMC

PBMCs (2.5 × 10^5^) were centrifuged at 350xg for 5 min at 4°C, the supernatant removed and the cells suspended in either Ad5-EGFP or Ad5F35-EGFP in serum-free RPMI and incubated for one hour at 37°C in a humidified atmosphere with 5% CO_2_. Complete RPMI 1640 was added to each sample and incubated for a further 24 hours at 37°C in a humidified atmosphere with 5% CO_2_. The cells were centrifuged at 350xg for 5 min at 4°C, suspended in 1 ml PBS and centrifuged at 350xg for 5 min at 4°C. The supernatant was removed and transduced cells identified by EGFP expression and flow cytometry, with different lymphocyte populations identified using antibodies to identify CD3+ (all T cells), CD4+CD3+ (CD4+ T cells), CD8+CD3+ (CD8+ cells) and CD56+CD3^neg^ (NK cells).

### Transduction of Primary NK cells

NK cells (2.5×10^5^) isolated by indirect selection from PBMC were centrifuged at 350xg for 5 min and washed once with PBS. The supernatant was removed, cells were suspended in serum-free RPMI and treated with Ad5-EGFP or Ad5F35-EGFP. Cells were incubated for one hour at 37°C in a humidified atmosphere with 5% CO_2_, 1 mL of complete RPMI 1640 medium was added and incubated at 37°C for a further 24 h in a humidified atmosphere with 5% CO_2_. The cells were collected by centrifugation, washed in PBS with centrifugation (350xg, 5 min) and suspended in PBS; transduction was analysed by EGFP expression and flow cytometry.

### Detection and quantitation of cell-surface CD46 molecules by flow cytometry and microsphere bead assays

Adherent cell lines were detached using protease-free Versene (Sigma-Aldrich) followed by addition of complete DMEM, centrifugation at 350xg for 5 min and resuspension in PBS plus 1% Bovine Serum Albumin (BSA). Suspension cells were centrifuged at 350xg for 5 min and suspended in PBS plus 1% BSA. Each sample for flow cytometry comprised 2.5×10^5^ cells. Cells were incubated with 1% (final concentration) mouse serum (for CD46) for 10 min on ice (to block non-specific immunoglobulin binding sites) followed by addition of PBS. Cells were collected by centrifugation (350xg, 5 min) and incubated with FITC-conjugated mouse anti-CD46 for 30 min on ice, washed twice in PBS with centrifugation (350xg, 5 min). Cells were washed twice in PBS with centrifugation (350xg, 5 min), analysed by flow cytometry on a FACSCalibur, (BD Bioscience, Wokingham, Berkshire, UK) or Attune (Thermo Fisher Scientific; Waltham, MA, USA) and the data processed using FlowJo software (Tree Star, Ashland, OR, USA).

For quantitation of cell-surface CD46 molecules, QSC microsphere beads were used, according to the manufacturer’s instructions (Bangs Laboratories, IN, USA). The geometric means of the QSC microsphere beads and cell samples were entered into the QuickCal analysis template provided by the manufacturer, generating a standard curve and readings of the number of cell-surface molecules present on cells.

### Determination of cell-surface area by cell labelling and Lattice LightSheet microscopy

Single cell suspensions of adherent epithelial cells were prepared as described above. One µl of stock CellTrace Yellow dye (Invitrogen, UK #C34573) was added to suspensions of NK or epithelial cells and incubated at 37°C for 20 minutes. Five volumes of complete medium was added to each tube and incubated for a further 5 minutes at room temperature. Cells were collected by centrifugation as described above. The labelled cell pellets were resuspended in complete medium and incubated at 37°C for 10 minutes. Aliquots (500 µl) of labelled cells were transferred to 35mm glass bottom dishes (Ibidi, Germany #C81218-200). Images were acquired using a Lattice Light sheet-7 microscope (Zeiss, Germany) and used to calculate cell-surface area using Imaris x64 V9.3.0 software. Single cells and their cell surface areas were calculated using both manual and automatic options. The most strongly labelled cells were chosen for the calculation of average cell surface areas using semiautomatic options. The average surface area (µm^2^) and the density of CD46 molecules per µm^2^ for each cell line was determined.

### Transcriptome and proteome data from public databases

Bulk RNAseq data for genes under test was downloaded from the functional annotation of the mammalian genome 5 (FANTOM5) dataset [24,25], available at: https://fantom.gsc.riken.jp/5/. This data includes quantitation of expression from multiple transcriptional start sites. This was converted to a single expression value for each gene by summing reads from the annotated promoters. Single cell (sc) RNAseq data was visualised and analysed using Single Cell Portal [26], available at: https://singlecell.broadinstitute.org/single_cell. Studies analysed via Single Cell Portal were from human lung [27] and human blood [28]. Human cell surface proteome data [29] was obtained via http://wlab.ethz.ch/cspa.

### Statistical Analysis

Statistical analysis was performed using Graphpad Prism software and Microsoft Excel. The statistical test used in each experiment is indicated in the figure legends.

## 3. Results

### Expression of the CD46, DSG2 and CXADR genes in human tissues and cell lines

The fibre from Ad35 binds to CD46 and replacement of the Ad5 fibre with that from Ad35 generates a chimaeric Ad capable of transducing CD46 expressing cells [14,30]. We previously demonstrated that Ad5F35 could transduce B, T and NK cell lines and the epithelial cell line HeLa [20]. This transduction of both haematopoietic and epithelial cell types reflects the widespread expression of CD46, in contrast to the more restricted expression of CAR and DSG2 [8,31]. However, to the best of our knowledge, an extensive comparison of expression of Ad entry receptor molecules across human cell types has not been reported. The ability to interrogate large, public-domain gene expression databases allowed us to analyse expression of CD46, CXADR (encoding CAR) and DSG2 (desmoglein 2) genes across multiple human tissues and cell types. Data was taken from the FANTOM consortium, in which mRNA from 889 different human tissues, primary cells and cell lines was analysed by quantitative capped analysis of gene expression (CAGE) [24,25]. This method determines the sequence of the 5’ end of mRNAs, allowing mapping of transcriptional start sites and quantitation of reads. Expression of CD46, CXADR and DSG2 genes was analysed, along with PTPRC (encoding CD45, a marker of haematopoietic lineage cells) and EPCAM and CDH1 (encoding epithelial cell adhesion molecule and E-cadherin respectively) as markers of epithelial cells. Display of gene expression across all 889 samples revealed that CD46 mRNA was widely expressed, whereas CXADR and DSG2 gene expression was more restricted (Fig. 1A and Supplementary Table 1). Comparison of expression of CD46, CXADR and DSG2 in primary lymphocytes and epithelial cells showed that CD46 mRNA expression levels were not significantly different between these tissue types, whereas CXADR and DSG2 were preferentially expressed by the epithelial cell types (Figure 1B). A corresponding analysis of protein expression was performed using data from the Cell Surface Protein Atlas, a mass spectrometry-based analysis of cell surface proteins across 47 human cell types [32]. Expression of CD46 was detected in 37 of the 47 cell types (including primary T cells and NK cells), in marked contrast to CAR expression, which was only detected in three of the 47 samples (Figure 1C and Supplementary Table 2). Furthermore, CD46 was detected in all of the haematopoietic-derived cells (along with CD45), whereas CAR was not expressed in any haematopoietic sample (Figure 1C). These results demonstrate that CD46 is widely expressed among human cell types, highlighting the potential of species B Ads and their derivatives such as Ad5F35 for gene delivery to a wide variety of cells and tissues. We confirmed these patterns of gene expression at the single cell level using data obtained from human lung and blood studies [27,28]. The results confirm that CXADR and DSG2 are preferentially expressed in epithelial tissue whereas CD46 is expressed by multiple haematopoietic cell types (including both lymphocyte and myeloid cell populations) as well as epithelial cells (Figure 2A, B). Overall, these results confirm the broad expression of the CD46 gene and protein and highlight its potential as a target for achieving Ad entry into many tissue types.

**Figure 1.**
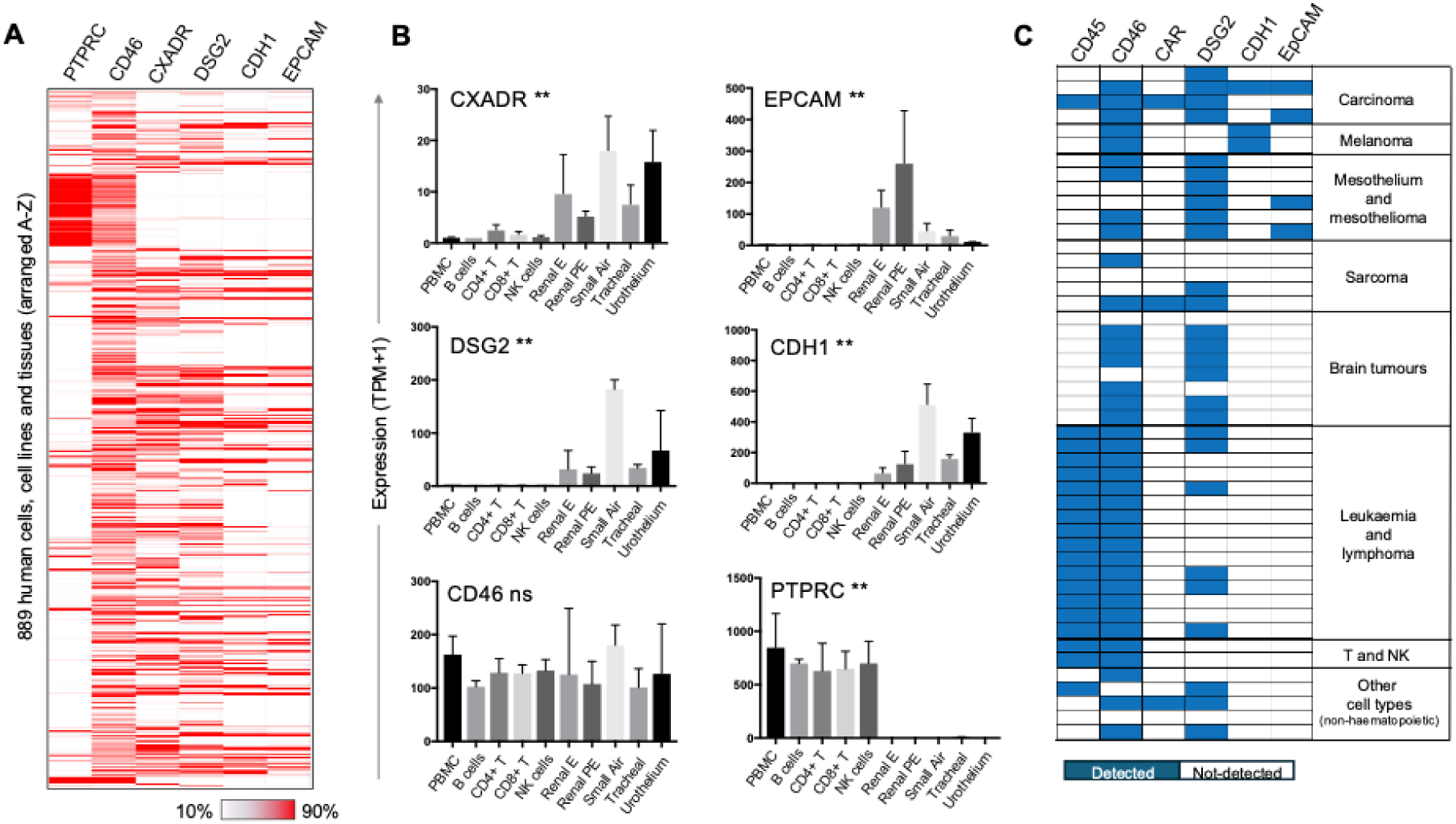
Expression of adenovirus entry receptors at the mRNA and protein level. **A)** Expression of genes for adenovirus entry receptors CD46, CXADR (encoding CAR) and DSG2 (desmoglein 2) across 889 human cells and tissues (arranged A-Z) using data from the FANTOM consortium [25]. Data for each gene was downloaded from https://fantom.gsc.riken.jp/5/sstar/Main_Page. Expression of PTPRC (encoding CD45) is shown as a marker of haematopoietic cells and both CDH1 (Epithelial cadherin/CD324) and EPCAM (Epithelial cell adhesion molecule/CD326) as markers of epithelial tissue. Expression levels are indicated according to the range in each column. The identity of the 889 tissues together with the expression values is provided in Supplementary Table 1. **B)** Expression of adenovirus entry receptor genes and marker genes in ten selected tissues (five haematopoietic and five epithelial) from the FANTOM consortium dataset. Expression levels across the indicated tissues were analysed using a Krustal Wallis test (**P<0.01, ns not significant). Tissues are peripheral blood mononuclear cells (PBMC), B cells, CD4+ T cells, CD8+ T cells, natural killer (NK) cells, renal epithelial (Renal E) cells, renal proximal tubular epithelial (Renal PE) cells, small airway epithelial (Small air) cells, tracheal epithelium (Tracheal) and urothelium. Number of samples per column is n=3. **C)** Expression of adenovirus entry receptors and marker molecules at the protein level as determined by mass spectrometry across 47 samples. The data is from the Cell Surface Protein Atlas [32], available at http://wlab.ethz.ch/cspa and shows whether the indicated protein was detected or undetected as indicated. A full list of the samples analysed is provided in Supplementary Figure 1.

**Figure 2.**
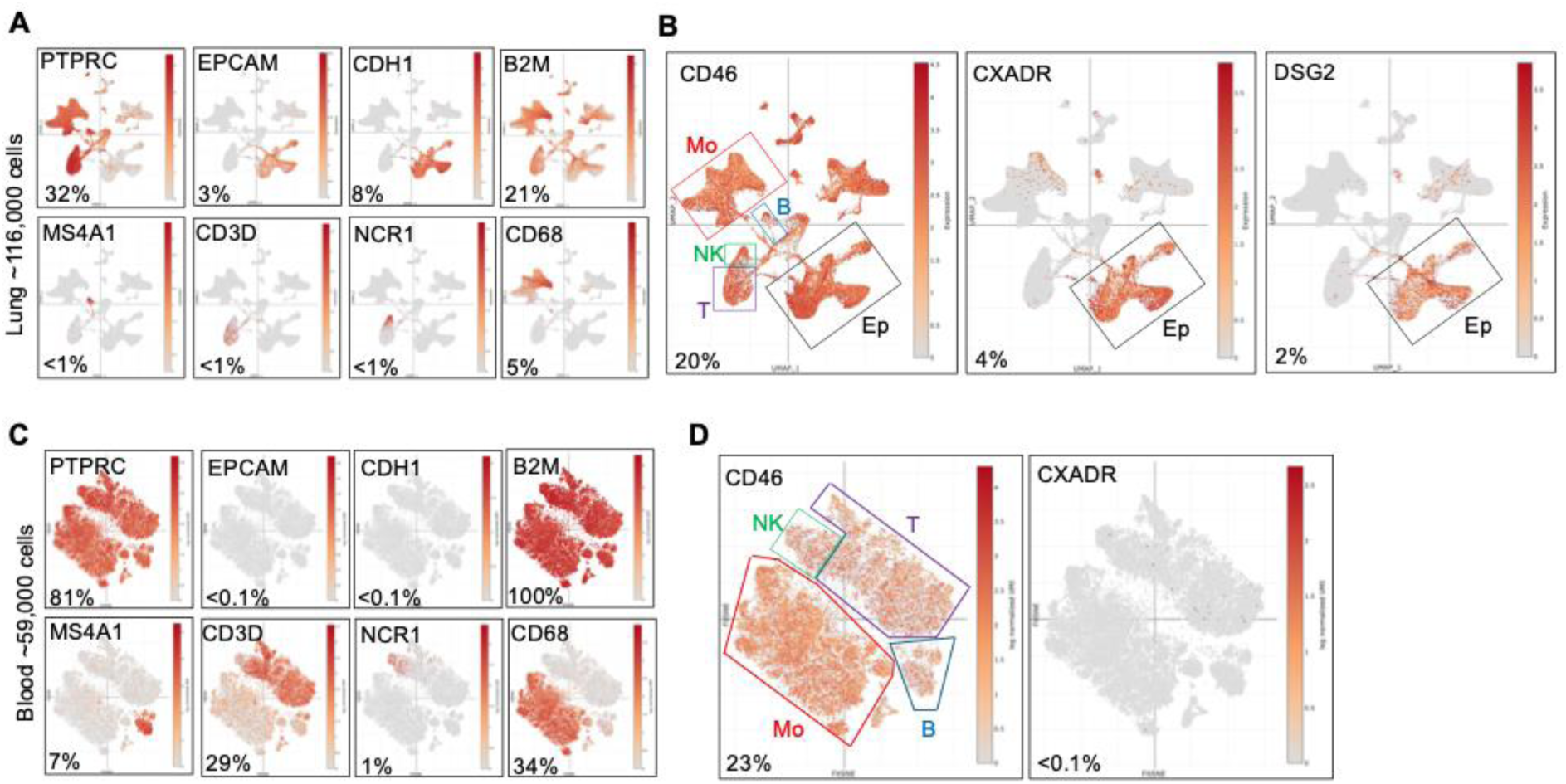
Expression of Adenovirus entry receptors at the single cell level. Expression of Adenovirus entry receptor genes (CD46, CXADR, DSG2) and lineage marker genes analysed at the single cell level. **A)** 116,000 lung cells [27] and **B)** in 59,000 blood cells [28]. Data was viewed and analysed using Single Cell Portal [26]; https://singlecell.broadinstitute.org and shows cell populations with expression of indicated genes superimposed. The percentages shown are the proportion of total cells expressing the named gene. Expression levels of DSG2 were not determined in the blood study. The marker genes used to identify major populations of cells are PTPRC (haematopoietic cells), EPCAM and CDH1 (epithelial cells), MS4A1 (B cells), CD3D (T cells), NCR1 (NK cells), CD68 (monocyte lineage cells) and B2M (housekeeping gene).

### Ad5F35 transduction of primary human NK cells

We previously showed that the human NK cell line NK92MI was refractory to transduction with Ad5-EGFP, but that transduction was possible with Ad5F35-EGFP. However, transduction of NK92MI by Ad5F35 required approximately ten times more virus particles per cell (ppc) compared to transduction of Hela cells. Furthermore, use of peripheral blood mononuclear cells (PBMC) demonstrated transduction of NK cells, T cells and B cells when high titres of Ad5F35 were used [20]. We explored the transduction of primary NK cells in more detail. Human primary NK cells were purified from peripheral blood mononuclear cells (PBMC) using magnetic, indirect immunoselection, generating a highly pure population of NK cells defined as CD56+CD3^neg^ cells (Supplementary Figure 1). Purified NK cells were transduced with an increasing MOI of Ad5F35-EGFP and expression of EGFP analysed 24h post transduction using flow cytometry. Similar to our previous results [20], efficient transduction of primary NK cells required 10-20 fold more Ad5F35 particles per cell than was required for A549 lung epithelial cells. In addition, the transduced NK cells showed no significant difference in viability compared to their untransduced counterparts (Supplementary Figure 1). It was possible that the high number of Ad5F35 virus particles used to transduce NK cells bypassed the requirement for the Ad35 fibre for entry. However, comparison of primary NK cell transduction by Ad5- or Ad5F35-EGFP showed that very high levels of Ad5-EGFP virus particles (up to an MOI of 2000) were unable to transduce primary NK cells and that NK cell transduction was indeed dependent on the Ad35 fibre protein (Figure 3A). The requirement for CD46 in the transduction of NK cells by Ad5F35 was tested with a blocking polyclonal anti-CD46 antibody. Primary NK cells from three donors and A549 cells were transduced with Ad5F35-EGFP in the presence of anti-CD46 blocking antibody (or control antibody). Treatment with the blocking antibody decreased the transduction of both primary NK cells and A549 cells, with the latter showing a statistically significant reduction (Figure 3B, C). This is consistent with a role for CD46 as an attachment/entry molecule for Ad5F35 in NK cells and epithelial cells.

**Figure 3.**
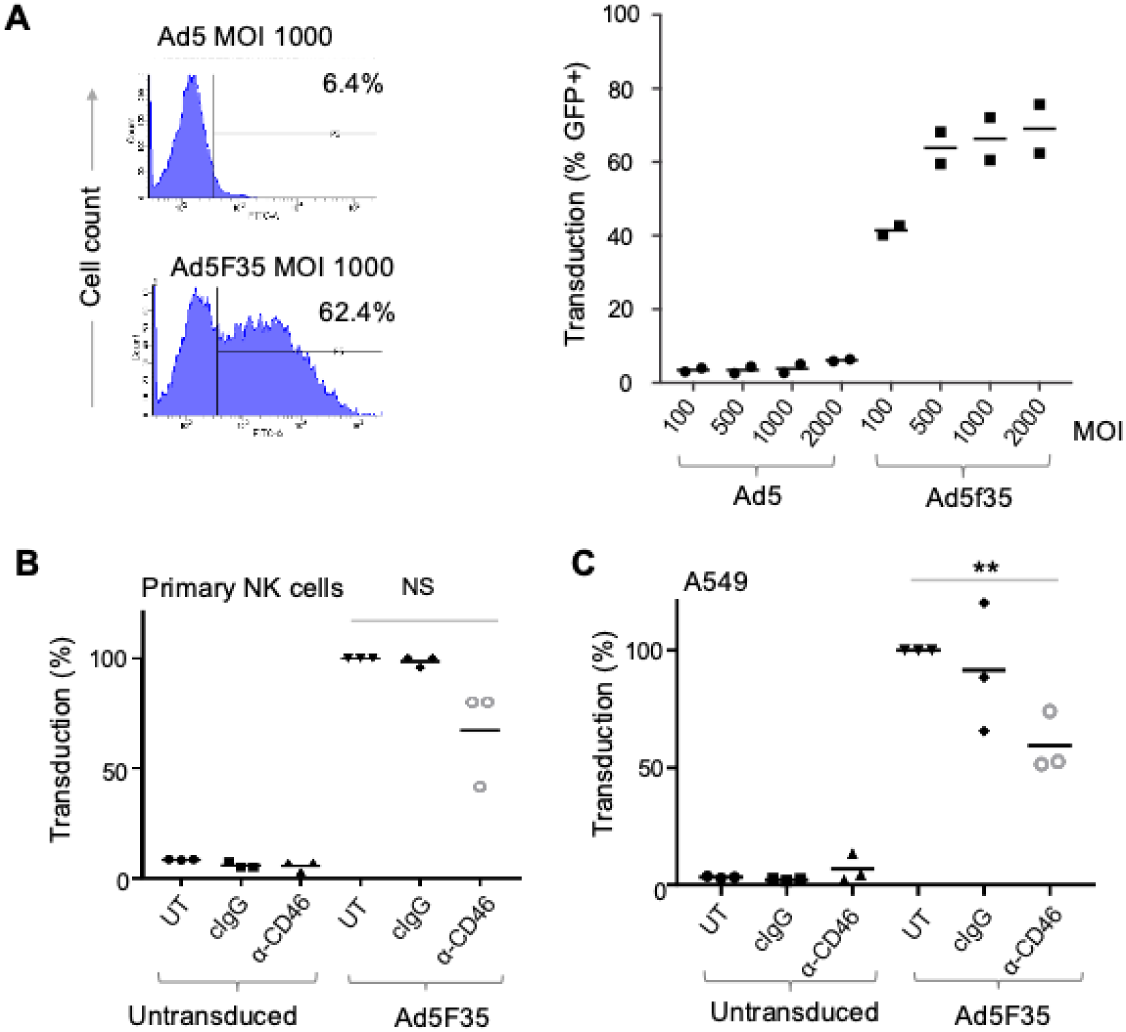
Ad5 and Ad5F35 transduction of primary NK cells and the A549 cell line. **A)** Left panel: Transduction of purified NK cells by Ad5-EE and Ad5F35-EGFP (MOI 1000), assessed by EGFP expression and flow cytometry; percentages show GFP+ cells. Right panel; transduction by an increasing MOI of Ad5-EGFP and Ad5F35-EGFP on purified NK cells showing data from two separate donors with mean values. **B)** and **C)** Transduction (assessed as percentage of EGFP expressing cells) of **B)** primary NK cells and **C)** A549 cells in the presence of a blocking antibody against CD46 (α-CD46), a matched control antibody (cIgG) or untreated cells (UT). Untransduced cells were analysed as controls. Transduction of A549 cells was performed using MOI of 0.3 in three separate experiments and EGFP expression analysed 24 hrs post-transduction. Transduction of NK cells used 3 separate donors and an MOI of 500; EGFP expression was analysed 24 hrs post-transduction. Data was analysed using a paired t test for NK cells (pairing data within donors) and an unpaired t test for the A549 cell line; **P<0.01, NS, not significant.

### The density of CD46 molecules at the cell surface

We hypothesised that the different amounts of Ad5F35 needed to efficiently transduce primary NK cells and A549 cells reflected the relative levels of cell-surface CD46 on each cell type. The number of CD46 molecules on A549 cells was compared to NK cells (and T cells) using the antibody binding capacity (ABC) approach. This compares the antibody-based fluorescence signal from stained cells with that obtained by staining of calibration beads of different sizes containing known numbers of antibody binding sites [33,34]. The fluorescence signal obtained following staining of the calibration beads with saturating levels of PE-conjugated anti-CD46 antibody (Figure 4A) was used to construct a standard curve of antibody binding capacity versus resultant mean fluorescence intensity (Supplementary Figure 2). Although bivalent, a single IgG molecule binds predominantly to a single antigen when present at saturating levels, indicating that the ABC is approximately equal to the number of antigens on the target cell [33,35]; PBMC were reacted with antibodies to detect NK cells (defined as CD56+CD3neg) or T cells (CD3+) and CD46 expression was analysed on these lymphocytes in parallel with A549 cells. Use of the fluorescence data in Figure 4A and interpolation of the standard curve showed an anti-CD46 ABC of approximately 21000 for NK cells and 30000 for T cells (Figure 4A). The fluorescence following staining of A549 cells with anti-CD46-PE exceeded the ABC of the highest density calibration bead, and hence an ABC of >625180 for these cells was recorded for A549 (Figure 4A). We were concerned that this very high ABC for anti-CD46 might reflect non-specific binding of this antibody to A549 cells. However, treatment of A549 cells with an siRNA targeting CD46 reduced the ABC by >95% compared to controls (Figure 4A). We then determined the ABC of NK cells, CD4+ T cells and CD8+ T cells from three separate donors using this technique. This demonstrated that cell surface CD46 levels demonstrated relatively little donor to donor variation and that NK cells had an anti-CD46 ABC of approximately 20000 and T cells were approximately two-fold higher (Figure 5B).

**Figure 4.**
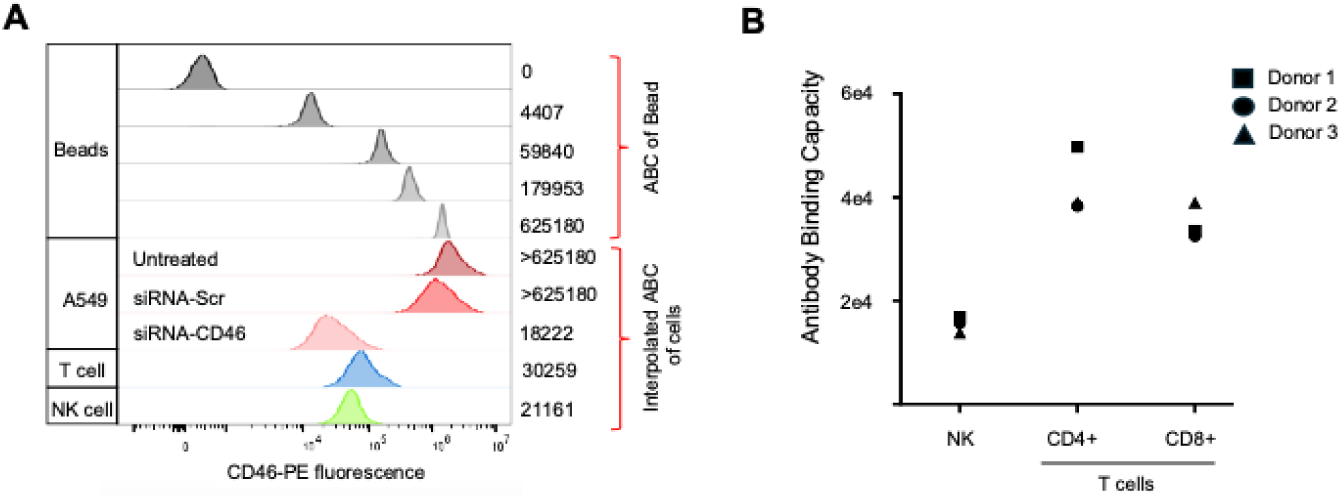
Anti-CD46 antibody binding capacity (ABC) of NK cells, T cells and the A549 cell line. **A)** Flow cytometry showing fluorescence of anti-CD46-PE binding to beads with defined antibody binding capacity (ABC of bead shown on right). This data was used to construct a standard curve of ABC versus fluorescence (shown in Supplementary Figure 2). Also shown is fluorescence of anti-CD46-PE staining of primary NK cells, T cells and A549 cells, with the interpolated ABC for this biding shown on the right. For A549 cells, staining with anti-CD46-PE was also performed on A549 transiently transfected with siRNA targeting CD46 (siRNA-CD46) or a control sequence (siRNA-Scr) to validate the very high levels of CD46 on these cells. **B)** ABC of NK cells, CD4+ T cells and CD8+ T cells for anti-CD46-PE using cells from three separate donors.

**Figure 5.**
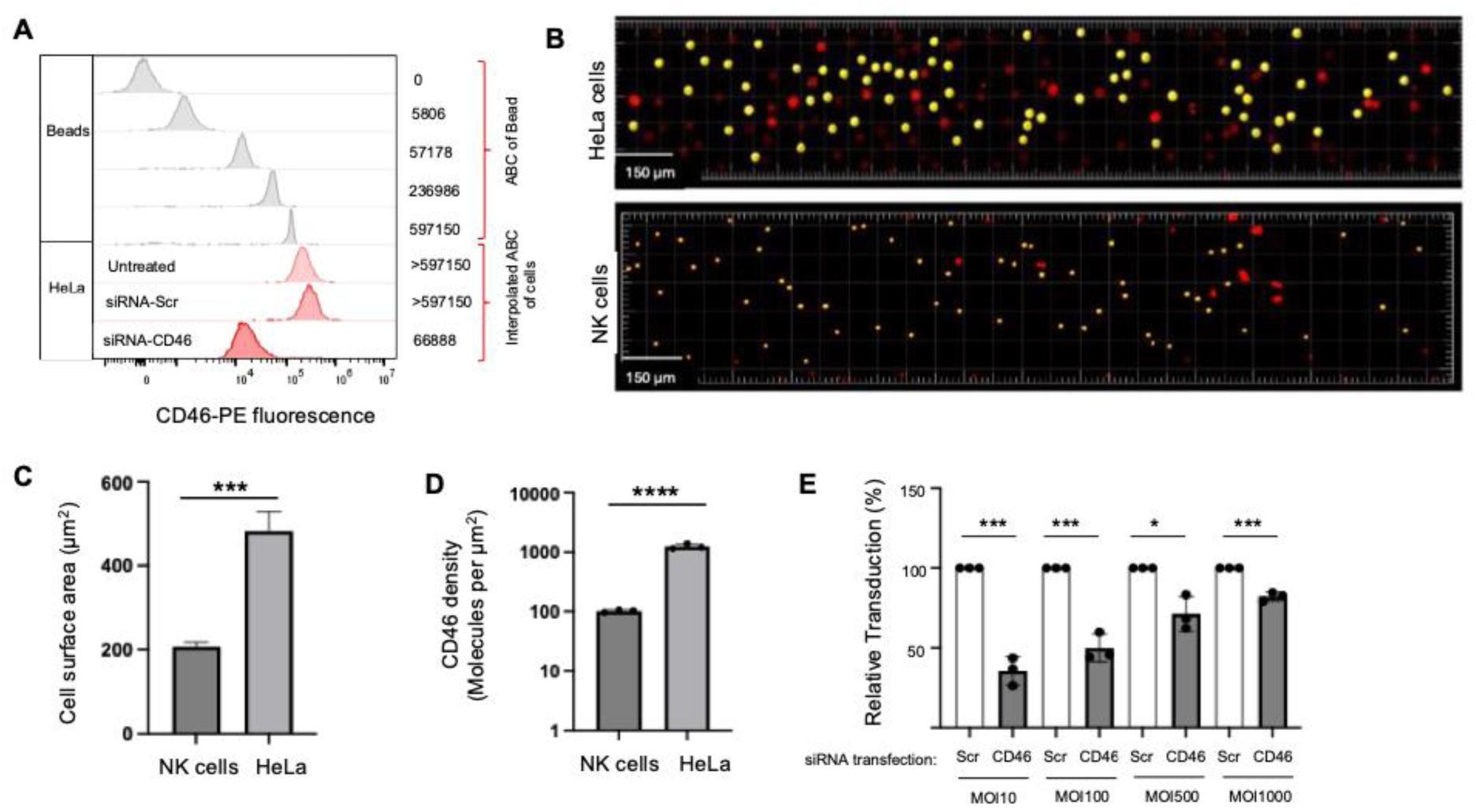
CD46 density on NK cells and HeLa cells. **A)** Flow cytometry showing fluorescence of anti-CD46-PE binding to beads with defined antibody binding capacity (ABC of bead shown on right) and fluorescence of anti-CD46-PE staining of untreated HeLa cells or HeLa cells transiently transfected with siRNA targeting CD46 (siRNA-CD46) or a control (siRNA-Scr) to validate high levels of CD46 on these cells. **B)** Light sheet microscopy of human NK cells and HeLa cells to determine cell size. Cells were labelled with CellTrace Yellow and imaged using a Zeiss Lattice LightSheet-7 microscope using a red filter; cells labelled in yellow were used in analysis. **C)** Cell surface area (µm^2^) of NK cells (n=268) and HeLa cells (n=370) as determined by light sheet microscopy. Data for each cell type is pooled from three separate experiments; in the case of NK cells this was performed using two separate donors (one donor analysed in one single experiment and one donor analysed in two separate experiments). **D)** Cell surface density of CD46 molecules on NK cells and HeLa cells (Molecules per µm^2^). This was calculated using the ABC values for CD46 on these cells and the mean cell size as determined by light sheet microscopy. **E)** Effect of CD46 density on Ad5F35-EGFP transduction of HeLa cells. HeLa cells were transiently transfected with siRNA-CD46 or Scr control and then transduced with Ad5F35-EGFP across the range of MOIs shown.

These results suggest that A549 cells express a minimum of 30-fold more CD46 molecules than NK cells. We performed this analysis on HeLa cells (Figure 5A). Similar to A549 cells, HeLa cells exceeded the ABC of the highest density calibration bead used and we therefore recorded an ABC of >597150; this was reduced by ∼90% following knockdown of CD46 expression using siRNA (Figure 5A). Like A549 cells, HeLa cells (ABC >597150) expressed approximately 30-fold more CD46 molecules per cell than NK cells (ABC 20000).

Lymphocytes, including NK cells, are smaller than epithelial cells. This prompted an evaluation of the density of CD46 molecules on NK cells and epithelial cells, since variation in cell density of CD46 might account for the increased numbers of Ad5F35-EGFP virus particles required to transduce NK cells compared to epithelial cells. We used Lattice LightSheet microscopy [36] to determine cell surface area. This method can be used to accurately image cells in suspension (essential for NK cells) using fluorescence-labelling for cell detection and measurement. Furthermore, the method exhibits reduced photobleaching, allowing for greater number of cells to be accurately sampled and analysed; Imaris software running on the Zeiss Lattice LightSheet-7 microscope facilitates cell size determination. Suspension cultures of each cell type were incubated with CellTrace Yellow dye to uniformly label the cells, generating labelled spheres suitable for comparison. The images clearly demonstrate the difference in size between the primary NK cells and HeLa cells in suspension (Figure 5B). Measurement of the surface area of the labelled sphere was taken to be equivalent to the cell surface. The surface area of the labelled spheres was quantitated, and a median area (in µm^2^) was calculated (Fig 5C). This demonstrated that HeLa cells (area ∼500 µm^2^) had a surface area approximately two and half times that of NK cells (area ∼200 µm^2^). Using the ABC data in Figures 4 and 5, the density of CD46 molecules (expressed as CD46 molecules per µm^2^) was found to be approximately 100 molecules of CD46/µm^2^ in primary NK cells and 2000 CD46/µm^2^ in HeLa cells (Figure 5D), a 20-fold difference.

### Ad5F35 transduction of HeLa cells with reduced CD46 density

We hypothesised that reducing CD46 density on HeLa cells would reduce Ad5F35 transduction, but that this could be overcome by increasing the MOI, mimicking the situation in NK cells. Transient siRNA transfection of HeLa cells reduced the cell-surface CD46 from an ABC of approximately >600000 in control siRNA-transfected cells (siRNA-Control) to an ABC of approximately 67000, representing greater than a 90% reduction in cell-surface CD46 (Figure 5A). Importantly, the 20-fold lower CD46 density on these siRNA-transfected cells more closely matches the density of CD46 on NK cells. Transduction of these siRNA-transfected HeLa cells with Ad5F35-EGFP showed that an increased MOI can compensate for the reduced density of CD46 at the cell surface (Figure 5E).

## 4. Discussion

Human adenoviruses (Ads) have attracted considerable recent interest as vaccine and gene therapy vectors, targeting diseases such as SARS-CoV2 coronavirus infection as vaccine vectors, as well as being developed for use as selective anti-cancer agents [37,38]. It is of fundamental importance to understand the Ad lifecycle in human cells so that specific Ad types can be chosen for applications, or that new genetic modifications can be introduced to create Ads with novel therapeutic potential [39]. In particular, the initial steps in Ad attachment to target cells and post-binding entry pathways are important in terms of target cell selection and efficiency of virus entry [40,41].

Cell surface CD46 molecules are potential targets for cancer therapy due to their elevated expression in cancer cells and tissues [42]. For example, therapeutic anti-CD46 antibodies conjugated to toxins have been developed which target myeloma cells with high CD46 expression while sparing non-cancerous blood cells with lower CD46 expression [43]. Cell-surface CD46 molecules are targeted by several pathogens, including certain human adenovirus (Ad) types (such as Ad35), human herpesvirus 6, certain measles virus (MV) strains and bacterial pathogens, including *Neisseria* and *Streptococcus* species [44,45] Coupled with the observed up-regulation of CD46 (and other complement regulatory proteins) in cancer cells and tissues compared to their normal counterparts, CD46 has emerged as an important target for selective biotherapy of cancers by virus vectors such as Ad35 and the Edmonston strain of MV [46–48].

We previously showed that Ad5F35, but not Ad5, had the ability to bind and enter lymphoid cell lines derived from NK (NK92MI) and T cells (Jurkat) as well as the corresponding primary cell populations in peripheral blood lymphocytes. However, a much greater MOI of Ad5F35-EGFP was required to transduce primary NK or T cell lines than HeLa epithelial cells [20]. These observations prompted this investigation into the distribution and density of cell-surface CD46 molecules used for Ad5F35 entry into NK and epithelial cells and their impact on transduction by this chimaeric virus.

Our analysis of transcriptomes of more than 800 human tissue samples for expression of CD46, CAR (CXADR) and DSG2 genes revealed that only CD46 exhibited widespread tissue and cell expression. CAR and DSG2 expression were more restricted to epithelial cell types and showed similar distribution to cadherin 1 (CDH1) and epithelial cell adhesion molecule (EPCAM) transcripts. Proteomic database analysis confirmed the broad distribution of CD46 in many cancer cell types as well as leukaemia, lymphoma and primary T and NK cells. Single cell transcriptomic data of CAR (CXADR) and CD46 expression in blood and lung tissues showed that CAR was expressed in epithelial cells of the lung, along with DSG2 and EPCAM, whereas CD46 was expressed in T, B, NK and monocyte cells and well as epithelial cell types. Furthermore, CXADR expression was barely detectable in haematopoietic cells. These studies extend, in an unbiased and high throughput manner, previous immunohistochemistry data on the expression of CAR protein in 100 cancer and 273 normal tissues, which revealed variable expression of CAR in cancer tissues and low, variable levels in normal samples [49]. Our expression analysis provided a basis for the study of the mechanism of entry into primary lymphoid cells by an Ad5 vector in which the virus fibre protein from Ad type 35 replaced the Ad5 fibre, termed Ad5F35.

The regulation of CD46 expression and localisation at the cell-surface is complex. Transcription of the human CD46 gene appears to be regulated by several transcription factors and signaling pathways that are increased in cancer versus normal cells. For example, binding motifs for STAT3 and wild-type p53 are present upstream of the start-site for CD46 transcription [50,51]. STAT3 is frequently activated in cancer cells and tissues, resulting in increased expression of cell-surface CD46, protecting tumour cells from complement-mediated lysis and contributing to immune evasion [52,53]. Therefore, Ad5F35 and other Ads that utilise CD46 as an attachment molecule may be particularly useful as anti-cancer virotherapies.

Transduction of primary NK cells with Ad5- or Ad5F35-EGFP highlighted the origin of the fibre protein as an essential vector component that mediated selective targeting of NK cells, since these cells were not targeted by Ad5-EGFP. The fibre is a trimeric structure consisting of a globular carboxy-terminal head or knob domain that interacts with cell-surface attachment molecules, a shaft consisting of a variable number of repeated peptide sequences and an amino-terminal domain that embeds the fibre in the pentameric penton base protein. It is tempting to speculate that different structures of the Ad5 and Ad35 head domains explain the different interactions between these viruses and their cell-surface attachment molecules (CAR and CD46 respectively). However, the length of the fibre shaft has also been shown to affect attachment of Ads to cells [54]. The Ad35 fibre shaft has six repeat sequences whereas Ad5 has 22. The flexibility (or lack thereof) of the fibre due to its length may be a factor in binding of Ad5F35 to the NK cell surface and needs further investigation. Although the Ad5 penton base is present and identical in both Ad5 and Ad5F35, the interaction of its RGD motif with cell-surface integrins might be different in the two viruses. Previous studies have identified a key role for integrins in binding Ad35 virus to the surface of human haematopoietic and epithelial cells [55,56]. While the major integrins utilised for Ad entry into epithelial cells are α_v_β_3_ and α_v_β_5_, in haematopoietic cells, α_M_β_2_ appears to be a major integrin utilised by the Ad5 penton base for entry, even in the absence of a cellular attachment molecule [57]. The spectrum of cell-surface integrins in the primary NK cells used in this study has not been determined, although previous studies have reported the presence of α_M_β_2_ in circulating NK cells (of the type used in this study) and at the synapse of NK and target cells [58]. The combination of the penton base: integrin interaction and the presence of the primary attachment molecule for Ad5F35, namely CD46, on NK cells may provide an explanation for their susceptibility to Ad5F35 transduction. In contrast, interaction between the penton base and integrin alone in Ad5 appears insufficient to mediate cell entry.

The relationship between levels of CD46 expression and cell entry of the Edmonston strain of MV or Ad5F35 has been previously studied in a panel consisting of 16 clones of Chinese Hamster Ovary (CHO) cells stably transfected with human CD46 [47]. The results showed an approximately linear relationship between the level of surface CD46 molecules (as judged by flow cytometry, in terms of percent EGFP-positive cells) and transduction by the Edmonston strain of MV engineered to express EGFP. Similar results were obtained here with Ad5F35-EGFP. A related study was performed, in which numbers of CD46 molecules per cell were determined in multiple myeloma patients using a quantitative bead assay of patient-derived CD138^+^ neoplastic plasma cells (PC) and non-neoplastic CD138^neg^ plasma cells (NPC) from the same patient. This revealed an approximately seven-fold increase in CD46 molecules in CD138^neg^ PC versus CD138^+^ NPC cells. There was also preferential killing of PC compared to NPC cells by MV in most patient plasma cells. This advances the notion that CD46 is up-regulated in cancer versus normal cells and suggests that CD46 is a differentially expressed marker for targeted cancer therapy, for example by viruses [48]. In our study, cell-surface CD46 in primary NK and T cells from PBMC and epithelial cells (HeLa and A549) was quantitated using a bead-based assay. Since lymphoid and epithelial cells are of differing sizes, cell-surface areas were measured using Lattice Light Sheet microscopy of uniformly fluorescently-labelled cells, allowing a correction for cell surface area and accurate determination of cell-surface density of CD46 in molecules per µm^2^. This analysis revealed a 20-fold difference in cell-surface density between NK and Hela epithelial cells.

Unlike NK cells, HeLa cells are readily transiently transfected with siRNA. Resultant levels of HeLa cell surface CD46 were approximately 5% of control levels when siRNA targeted, reducing HeLa cell surface CD46 to a level similar to that found on NK and T cells. This makes CD46-low HeLa cells a model in which the impact of low CD46 density on Ad5F35 transduction can be comparatively assessed. Transduction of such low CD46-low cells with increasing numbers of Ad5F35 particles yielded approximately similar levels of transduction to that in NK cells. For example, approximately 40% of primary, untreated NK cells were transduced at an MOI of 100 compared with approximately 50% of HeLa cells that had been treated with CD46 siRNA and transduced at an MOI of 100. This demonstrates that the cell surface density of CD46 molecules is a major factor in determining the level of Ad5F35 entry into cells but that Ad5F35 transduction efficiency into cell types with varying CD46 levels can be optimised by alteration in the multiplicity of infection.

## 5. Authorship Contribution

TB, EA and SLD performed and analysed experimental work; MK prepared and tested viruses; EW supervised and analysed flow cytometry; RH supervised imaging and microscopy; GEB and GPC devised the concept and experimental strategies, wrote the manuscript and obtained funding for the work.

## 6. Acknowledgements

We thank Sally Boxall, Victoria Easton, Andy Berry and Liz Straszynski for help with flow cytometry analyses. We are grateful to our colleagues who provided advice and comments on this work.

