## Supplementary Data for "Targeting of immune cells by human adenoviruses: CD46 receptor density influences entry of chimaeric Ad5F35 into natural killer cells"

**Supplementary Figure 1:**

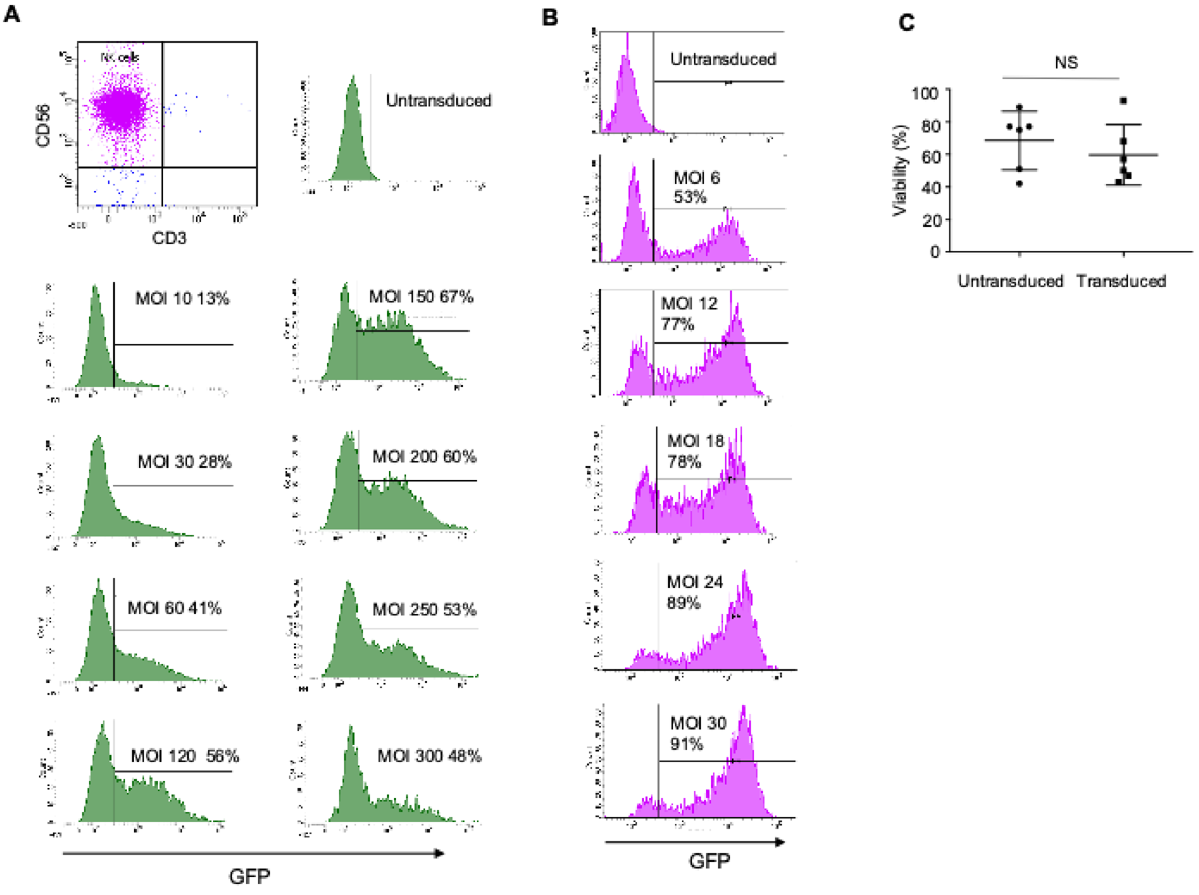


**Transduction of NK cells and A549 cells with an increasing MOI of Ad5F35-GFP.**

**A)** Primary human NK cells were isolated from peripheral blood mononuclear cells (PBMC) using indirect selection (Miltenyi Biotech) and the purity of the resultant cells tested by expression of CD56 and CD3; human NK cells are defined as CD56+CD3neg cells. These NK cells were transduced with Ad5F35-EGFP at a range of MOIs and EGFP expression used to identify transduced cells 24 hrs later (percentage transduced at each MOI indicated).

**B)** Transduction of A549 cells with Ad5F35-EGFP across a range of MOIs. Transduction was determined by EGFP expression 24 hrs post-transduction. Percentage of cells transduced at each MOI is indicated.

**C)** Viability of NK cells transduced by Ad5F35-EGFP. NK cells were transduced with Ad5F35-EGFP, stained with a dead cell discriminator and the percentage of live cells analysed in both the untransduced (non-EGFP expressing) and transduced (EGFP expressing) gates. This experiment was performed in six separate donors. Viability of untransduced and transduced cells within individual donors was not significantly (NS) different using a paired t test.

**Supplementary Figure 2:**

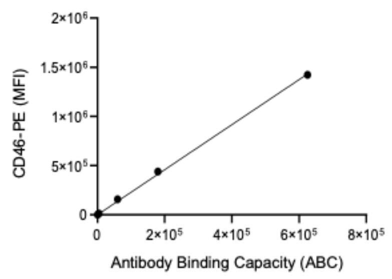


Standard curve of anti-CD46-PE fluorescence using beads with defined ABC, as shown in Figure 4 of the main text. The standard curve was constructed using software provide by the manufacturer.

**Supplementary Tables:**

Supplementary Tables 1 and 2 provided as separate Excel files

**Supplementary Table 1:**

Dataset for Figures 1A and 1B. The 889 human cell types and tissues analysed by the FANTOM consortium [25] are listed from A-Z in column A with expression values for the six indicated genes shown in columns B-G. Expression values are coloured in each column according to whether the expression value is between 10% and 90% of the maximum expression level in that column. The data is based on capped analyses of gene expression (CAGE) which quantitates expression from multiple start sites for each gene. Expression values shown here are total expression in each sample. This was calculated by adding expression values obtained from each start site (promoter p1, p2 etc) annotated in the FANTOM dataset. Data was downloaded from; https://fantom.gsc.riken.jp/5/sstar/Main_Page

**Supplementary Table 2:**

Extended version of Figure 1C showing adenovirus entry receptors and marker molecules and whether they were detected (blue) or not (white) in the cell types indicated. Source data is from the Cell Surface Protein Atlas [32], available at http://wlab.ethz.ch/cspa.
